# Tasks and responsibilities in physical activity promotion of older patients during hospitalization: A nurse perspective

**DOI:** 10.1101/482976

**Authors:** Kira Scheerman, Joram W. Mesters, Jay N. Borger, Carel G. M. Meskers, Andrea B. Maier

## Abstract

**Objective:** To investigate how nurses perceive tasks and responsibilities in promoting physical activity during hospitalization of older patients and which factors are of influence.

**Design:** Observational cohort study

**Setting and participants:** One hundred and eight nurse students, nurses and nurse supervisors employed by an academic Dutch teaching hospital participated in a questionnaire survey and 51 nurses took part in a subsequent in-depth interview.

**Measures:** Data were collected on tasks and responsibilities in physical activity promotion and their influencing factors as perceived by nurses. Descriptive statistics were used to analyze the data from the questionnaire survey and a deductive approach with directed content analysis was used for the data from the interviews.

**Results:** Nurses perceived to have a dominant role in physical activity promotion of older patients during hospitalization. Ninety percent of the nurses stated to be responsible for physical activity promotion and 32 percent stated to be satisfied with the actual level of physical activity of their patients. Influencing factors were low patient motivation, high workload causing priority shifts of tasks and the role of physicians.

**Conclusion:** Although their perceived dominant role in physical activity promotion, nurses identified a number of barriers interfering with actual level of physical activity. Improvement strategies should involve physicians, patients and carers.

## Introduction

More than one third of all hospital discharges in the US affects patients 65 years or older (Data from Healthcare Cost and Utilization Project database 2013). Older hospitalized patients spend most of the time lying in bed (1–5) while physical inactivity in this group is associated with functional decline (6, 7), readmissions (8), nursing home admissions and death (6).

The level of physical activity during hospitalization has been indicated as a modifiable risk factor for complications related to hospitalization (9). Patient related, organizational and environmental factors have been shown to contribute to physical activity promotion of hospitalized older patients (10–13). Nurses are a vital actor due to the high amount of patient contact hours. Based on the education and professional profile, nurses are expected to signal risks and perform tasks to promote physical activity during hospitalization (14–18). However, it was shown that nurses infrequently initiate physical activity during hospitalization (19, 20), whereas evidence on the perception of nurses on their role in physical activity promotion is scarce (21).

The aim of the study was to investigate how nurses perceive tasks and responsibilities in promoting physical activity and to gain insight in what factors influence physical activity promotion by nurses in older patients during hospitalization.

## Methods

### Study design

This study encompassed both questionnaire surveys (March to July 2016) and in-depth interviews (June to August 2017). The study was conducted at an academic teaching hospital in The Netherlands and was approved by the medical ethical committee of Amsterdam UMC. All enrolled nurses provided written informed consent.

### Participants

Nurses were eligible when they were 18 years and older, were working on wards providing care to patients 70 years and older and had provided care to at least one patient of 70 years or older in the previous month. Nurses working on the medium or intensive care unit were excluded. All wards, with exception of genecology and pulmonology, were included for the questionnaire component. Included wards for the interview component were: internal medicine, traumatology, oncological surgery and a combined ward of vascular surgery, nephrology and urology. Nurses were selected at random from staff lists using a numerical lot drawing. Participation in both the questionnaire as well as interview component was allowed. The questionnaire component addressed nurse students, nurses and nurse supervisors. For the interview component only nurses were included.

## Procedure

### Questionnaire component

A questionnaire was developed to investigate tasks and responsibilities in physical activity promotion and factors influencing physical activity promotion during hospitalization of older patients as perceived by nurses. Questions were compiled by interviewing three nurses after they filled in the questionnaire and tested on feasibility and nurses’ interpretation. Selected nurses were approached by the head of the ward or by a researcher (JB). Questionnaire completion took approximately 20 minutes.

The following data were collected: age, gender, educational level, work experience, education on physical activity promotion, self-perceived motivation to promote physical activity, responsibilities in physical activity promotion, self-perceived level of knowledge of physical activity promotion, satisfaction of physical activity promotion of the ward, satisfaction of physical activity promotion of physicians and satisfaction of level of physical activity of the patient. Nurses were asked if they considered daily activities (e.g. teeth brushing at the sink, walking towards the toilet), unsupervised additional physical activity (e.g. stretch and gait exercises), additional physical activity supervised by a nurse and additional physical activity supervised by other health care professionals as physical activity during hospitalization and whether they promote daily activities, additional physical activity or consulted other health care professionals. Furthermore, nurses were asked to score the importance of 34 factors in physical activity promotion of older patients during hospitalization using a Likert-scale from 1 (totally not important) to 5 (very important) and to indicate their most important factor. Factors were selected based on literature and were categorized in characteristics of the professional, patient, organization or intervention, and social factors (22–30).

### Interview component

Targeted semi-structured interviews were conducted to gain in-depth information on nurses’ perspective on task and responsibilities of different actors in physical activity promotion and factors influencing physical activity promotion. The interview design was evaluated on feasibility and interpretation of the questions during a pilot interview with two nurses and, after adjustments, re-evaluated in a second pilot interview with two different nurses. Selected nurses were approached by the head of the ward or by the researcher (JM). Interviews were held in private rooms and had a duration of 30 to 50 minutes. The interviews were held by one researcher (JM), the first ten interviews were supervised by a second researcher (KS). The interviews were audio recorded and notes were made.

A patient case was provided at the start of the interview, describing a 82 year old woman with urosepsis, who had high fever and showed signs of delirium at admission, progressively got better at days two of admission and almost completely recuperated at day five of admission. Nurses were asked to define physical activity during hospitalization, to score the importance of physical activity promotion on a VAS-score and to describe how satisfied they were with the level of physical activity promotion during hospitalization. Tasks and responsibilities of nurses, physiotherapists, occupational therapists, physicians, patients, and carers were further explored with questions on the nurses’ perspective on responsibilities in signaling and performing different physical activity promotion tasks; transfer from bed to chair, activities of daily living, supervised and unsupervised additional physical activity. Nurses were also asked to motivate which actor they thought to have final responsibility of execution of these tasks and, subsequently, if these responsibilities would change when a patient would be able to, but not performing physical activity during hospitalization. Factors influencing physical activity promotion were discussed when nurses named factors during the interview and explicitly at the end of the interview supported with an overview of the 34 factors used in the questionnaire.

## Data processing and analysis

### Questionnaire component

Statistics were performed using IBM SPSS statistics 22.0. Data were expressed as number and percentages. Factors were considered important when Likert scores exceeded four. Fishers exact test was used to analyze differences between nurse students, nurses and nurse supervisors.

### Interview component

The interviews were fully transcribed and data on nurse characteristics, importance of physical activity promotion, tasks and responsibilities were analyzed quantitatively and qualitatively. A deductive approach with directed content analysis was used (31, 32) to analyze the qualitative data. Interview transcripts were read through and initial coding with pre-determined codes based on the topics of the interview were added to the correlating text. Open coding was used to refine labels. Atlas.ti 8.0 was used in the qualitative coding process. Discussions on interpreting data took place between two researchers (JM and KS) and all codes and data were verified by a second researcher (KS).

## Results

All 108 selected (student) nurses participated in the questionnaire component of the study. In the interview component, three nurses refused to participate resulting in a total of 51 interviews, see Table 1 for the characteristics of the participating nurses. Eighty-six percent of the nurses was female and the nurses in the questionnaire and interview component had a comparable median age (32 vs. 31 years) and level of work experience (6 vs. 7 years). In the questionnaire component, more nurses had a high educational level compared to the interview component (57.6% vs. 47.1%). A minority of nurse students (38.5%), nurses (20.0%) and nurse supervisors (30.0%) received education on physical activity promotion in the prior year.

**Table 1:** Characteristics of the study population, stratified by nurse student, nurse and nurse supervisor (questionnaire component, n=108 and interview component, n=51).

|  | Questionnaire |  |  | Interview |
| --- | --- | --- | --- | --- |
|  | Nurse student<br>n=13 | Nurse<br>n=85 | Nurse supervisor<br>n=10 | Nurse<br>n=51 |
| Age, y, median [IQR] | 25 [23.0-29.0] | 32 [25.0-51.0] | 49 [42.8-54.5] | 31 [26.0-45.0] |
| Female, n (%) | 11 (84.6) | 73 (85.9) | 10 (100) | 44 (86.3) |
| Educational level high<sup>A</sup>, n (%) | 7 (53.8) | 49 (57.6) | 10 (100) | 24 (47.1) |
| Work experience nurse, y, median [IQR] | N/A | 6 [3.0 – 20.0] | 25 [19.3– 30.3] | 7 [2.5- 18.0] |
| Work experience nurse supervisor,<br>y, median [IQR] | N/A | N/A | 7 [5.5 – 10.0] | N/A |
| Education about PA promotion last year, yes, n (%) | 5 (38.5) | 17 (20.0) | 3 (30.0) | N/a |

A: High educational level defined as applied university and higher; N/A: not applicable; N/a: not available; PA: physical activity.

### Tasks and responsibilities in physical activity promotion

The questionnaire component revealed that nurses feel responsible (89.4%) to promote physical activity during hospitalization. As presented in Table 2, nurses promote daily activities (95.3%) and consult other health care professionals to promote physical activity (90.6%). Seventy-eight percent of the nurses stated to actively promote additional physical activity like stretch and gait exercises. Only one nurse reported to not promote physical activity at all. No differences were observed between nurse students, nurses and nurse supervisors. Eighty-seven percent of the respondents stated that physiotherapists, physicians or the patients had responsibilities in physical activity promotion. Occupational therapists (13%) and carers (22%) were also mentioned to be responsible.

**Table 2:** Perception of physical activity promotion during hospitalization and methods to promote physical activity, stratified by nurse student, nurse and nurse supervisor (questionnaire component, n=108).

|  | Nurse student<br>n=13 | Nurse<br>n=85 | Nurse supervisor<br>n=10 |
| --- | --- | --- | --- |
| **Perception of physical activity promotion, n (%)** |  |  |  |
| Motivated to promote PA, yes | 12 (92.3) | 83 (97.6) | 10 (100) |
| Responsible to promote PA, yes | 13 (100) | 76 (89.4) | 10 (100) |
| Self-perceived level of knowledge about PA promotion, sufficient | 9 (69.2)* | 77 (90.6) | 8 (80.0) |
| PA promotion on ward, present | 8 (61.5) | 45 (52.9) | 6 (60.0) |
| Satisfied with PA promotion on ward, yes | 9 (69.2) | 56 (65.9) | 7 (70.0) |
| Satisfied with PA promotion by physicians, yes | 5 (38.5) | 41 (48.2) | 6 (60.0) |
| Satisfied with level of PA of patient, yes | 4 (30.8) | 27 (31.8) | 4 (40.0) |
| **Definition of physical activity during hospitalization, n (%)** |  |  |  |
| Daily activities | 12 (92.3) | 68 (80.0) | 8 (80.0) |
| Unsupervised additional PA of patient | 10 (76.6) | 66 (77.6) | 8 (80.0) |
| Supervised additional PA with nurse | 11 (84.6) | 70 (82.4) | 8 (80.0) |
| Supervised additional PA with other health care professionals | 12 (93.2) | 68 (80.0) | 8 (80.0) |
| **Methods of physical activity promotion during hospitalization, n (%)** |  |  |  |
| Promote daily activities | 12 (92.3) | 81 (95.3) | 10 (100) |
| Promote additional PA (e.g. stretch and gait exercises) | 9 (69.2) | 66 (77.6) | 8 (80.0) |
| Consult other health care professionals to promote PA | 11 (84.6) | 77 (90.6) | 10 (100) |
| I do not promote PA | 0 (0) | 1 (1.2) | 0 (0) |

PA; physical activity; \*: Difference between nurse student, nurse and nurse supervisor (P<0.05).

During the interviews, nurses stated to be responsible for signaling and performing physical activity promotion tasks and had final responsibility for transfers from bed to chair and promotion of daily activities. Nurses indicated that physiotherapists have a greater responsibility of supervised additional physical activity and it is the patients responsibility to do unsupervised additional physical activity. The majority of the nurses stated that patients’ responsibilities increase when patients become more independent during hospital admission and that they would motivate patients to be physically active by providing information on consequences of physical inactivity and discussing the reasons for the patients’ physical inactivity. The tasks and responsibilities of the physician were described as to determine the patient’s ability to perform different levels of physical activity and to motivate patients when they refuse to perform physical activity. The different tasks and responsibilities in physical activity promotion of all actors according to nurses are visualized in Fig 1.

**Fig 1:**
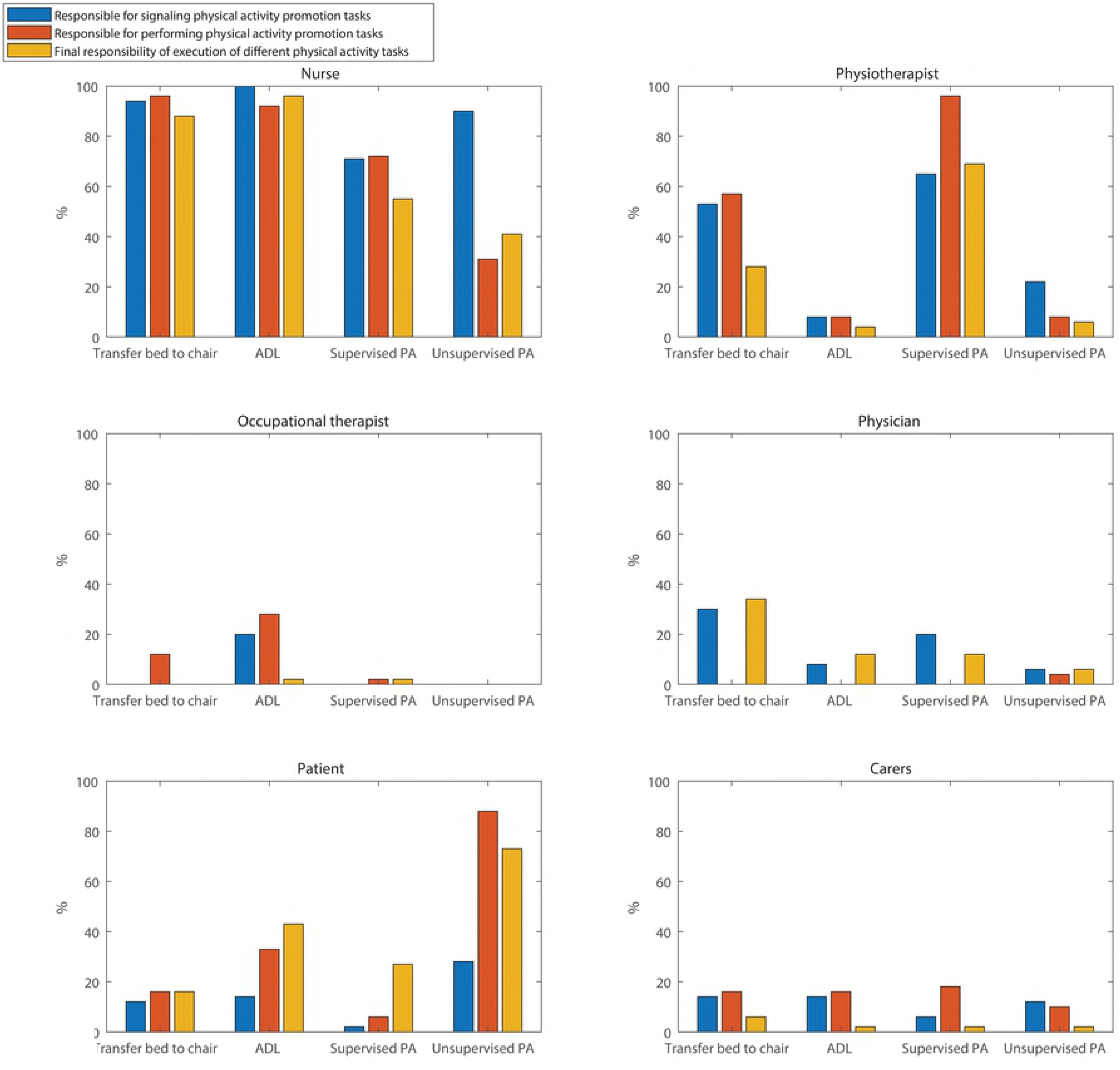
Tasks and responsibilities of different actors in physical activity promotion during hospitalization according to nurses (%) (interview component, n=51). PA: Physical activity

### Factors influencing physical activity promotion

Sixty-six percent of nurses of the questionnaire component were satisfied with physical activity promotion on their ward, 48% were satisfied with physical activity promotion by physicians, and 32% were satisfied with the actual level of physical activity of the patients during hospitalization. An overview of the importance of factors influencing physical activity promotion during hospitalization is provided in Table 3. Differences in importance between nurse students, nurses and nurse supervisors were observed for the factors ‘Fear of falling during PA promotion’, ‘Availability of protocol’ and ‘Opinion towards PA promotion of colleagues’.

**Table 3:**
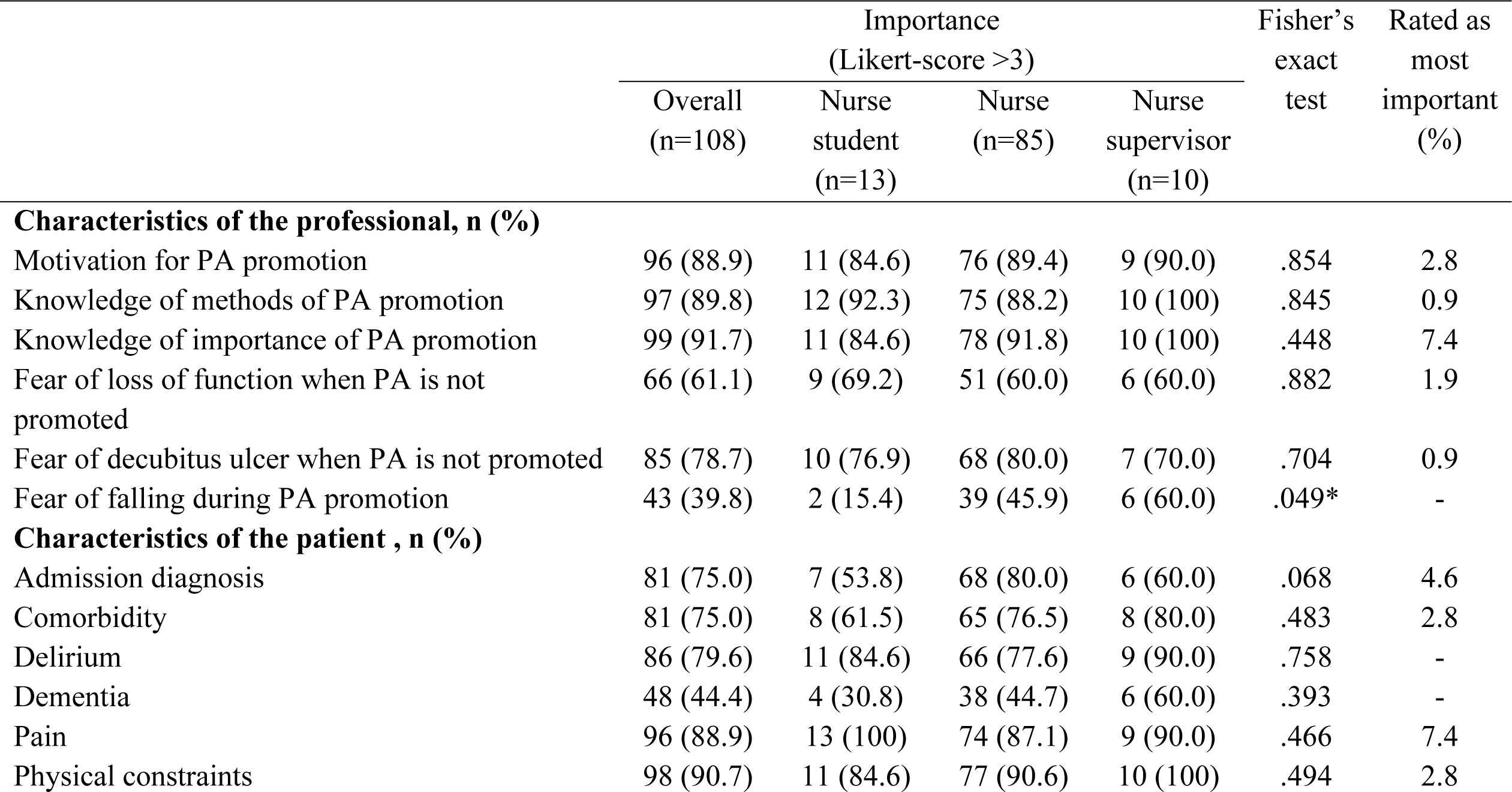
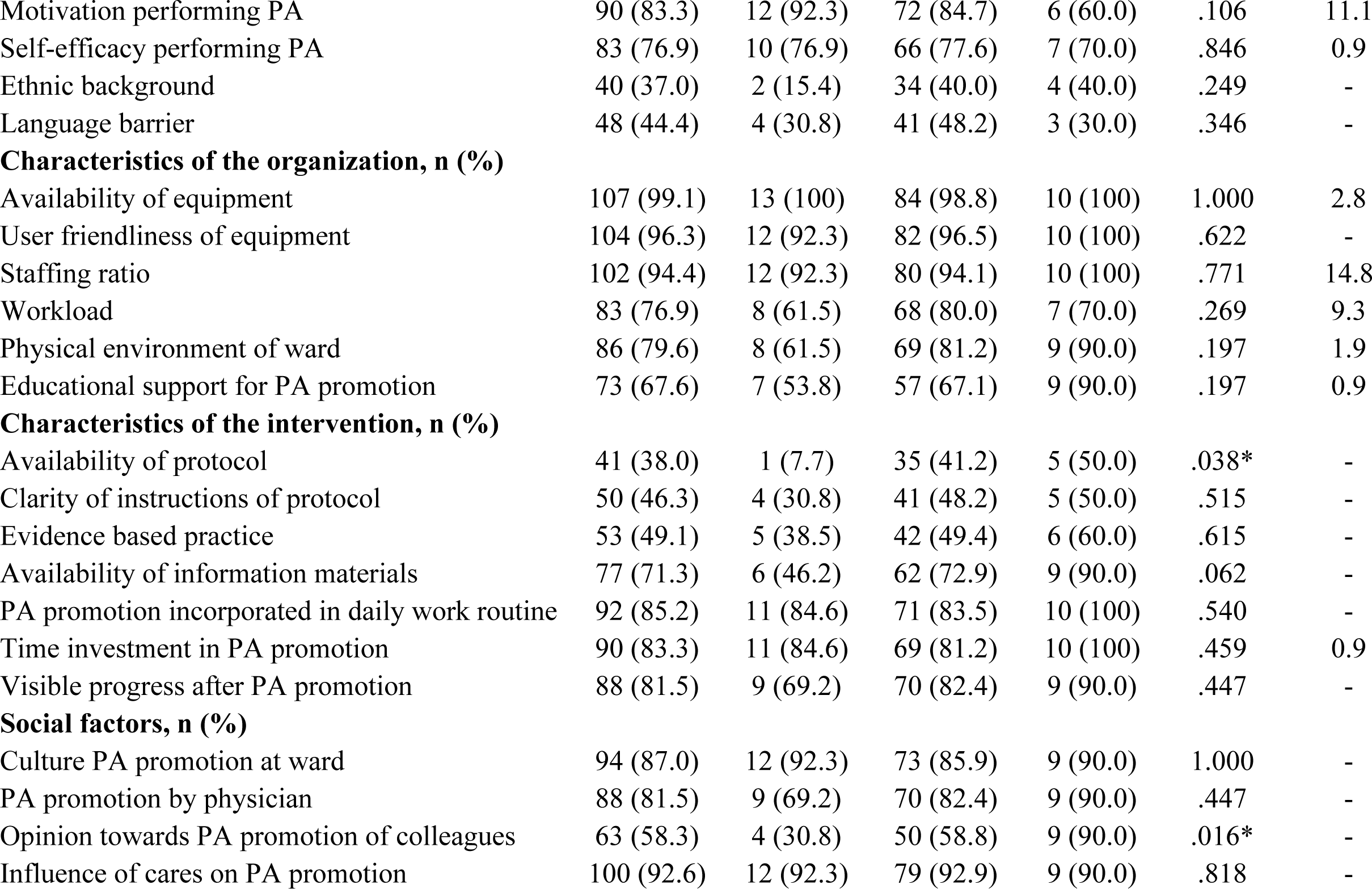
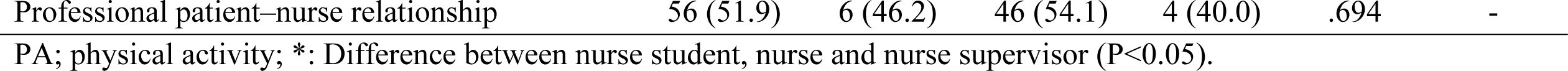
Importance of factors (Likert-score >3) influencing physical activity promotion in older patients during hospitalization, stratified by nurse student, nurse and nurse supervisor (questionnaire component, n=108).

### Characteristics of the professional

Almost all nurses scored knowledge of methods of physical activity promotion (88.2%), knowledge of importance of physical activity promotion (91.8%) and nurse’ motivation (89.4%) as important factors influencing physical activity promotion (Table 3). Nurses perceived their level of knowledge of physical activity promotion as sufficient (90.6%) in contrast to 69.2% of the nurse students (Table 2). Nurses were motivated to promote physical activity (97.6%) and considered daily activities (80.0%) and additional physical activity supervised by a nurse (82.4%) or other health care professional (80.0%) as physical activity during hospitalization (Table 2). Unsupervised additional physical activity was not seen as physical activity during hospitalization by 22.4% of the nurses. No differences were observed between the nurse students, nurses and nurse supervisors.

During the interviews, nurses described physical activity during hospitalization as: “out of bed”, “walking”, “movement in bed” and “sitting in chair”. Nurses scored the importance of physical activity during hospitalization with a median VAS score of 8.9 [5.5-10.0]. Most frequently named motivations were preventing risk of complications (e.g. decubitus and pneumonia), loss of muscle mass and slower recovery. Long term adverse outcomes like functional loss and regaining self-reliance were named less frequent compared with more immediate noticeable adverse outcomes. Nurses stated they were not always motivated to promote physical activity, because they were empathic towards patients facilitating comfort rather than physical activity. Also, the need for physical activity promotion in older patients was questioned. Nurses stated to understand the low motivation of older patients when being sick, had a bad night sleep or being tired after a day with multiple examinations.

### Characteristics of the patient

Physical constraints (90.6%), pain (87.1%) and motivation of the patient (84.7%) were identified as important factors in physical activity promotion (Table 3). The factor ‘motivation performing physical activity’ was scored as most important by 11.1% of the nurses.

During the interviews, patient motivation was stated as a barrier by 65% of nurses. Tiredness, pain, lines like catheters and drip-lines, their previous inactivity at home and the belief it is uncommon to be physically active at an older age were explanations for a low motivation of patients. In addition, the patient’ perception of the hospital admission (47%) as a place to rest, be sick and were it is justified to be physically inactive was identified as factor influencing the patient’ level of physical activity during hospitalization. Seventy-three percent of the nurses suggested hospitals to focus on increasing patient awareness on importance of physical activity, for example using a flyer or video.

### Characteristics of the organization

Sufficient staffing ratio (94.1%) and availability of equipment (98.8%) were identified as important factors in physical activity promotion (Table 3). The factor ‘staffing ratio’ was scored as most important by 14.8% of the nurses.

Nurses in the interview group stated that in case of a high number of complex patients (e.g. patients with delirium, physical impairments or multiple drip lines), workload and staffing ratio become barriers. Priorities shifts when nurses experienced a low staff ratio and a high workload; physical activity was regarded as one of the first activities to be dropped and nurses stated that concessions were made in their physical activity promotion (e.g. use of bed urinal). Nurses suggested hospitals to invest in more staff (physiotherapists and medical students), equipment and adjustment of patient rooms to make them more attractive for physical activity. A living room, walking routes and activity counseling were other suggestions to increase physical activity of the older patients during hospitalization.

### Characteristics of the intervention and social factors

Social factors like influence of carers (92.9%), culture of physical activity promotion on ward (85.9%), and physical activity promotion by the physician (82.4%) were indicated as important factors in physical activity promotion. Availability (41.2%) and clarity (48.2%) of a protocol regarding physical activity promotion were scored less frequently as important factor in the questionnaire component (Table 3).

## Discussion

Nurses perceive to have a dominant role in physical activity promotion and feel responsible, however, they were not satisfied with the actual level of physical activity of older patients during hospitalization. Low patient motivation and priority shifts of tasks due to high workload were indicated as barriers. In addition, the role of physicians was indicated to be important to influence physical activity promotion behavior.

### Tasks and responsibilities in physical activity promotion

This study indicated that nurses have to adopt various roles in physical activity promotion during older patients’ hospital admissions. Besides signaling and supporting physical activity promotion tasks and consulting other health care professional, nurses have an important role in motivating patients. Motivating patients and supporting self-management become more prominent in nursing (33, 34). However, nurse activities in physical activity promotion during hospitalization seem to target prevention of potential harm more than supporting rehabilitation goals (35). The reserved attitude of a part of the nurses in our study regarding physical activity of older patients during hospitalization and unsupervised physical activity suggests that the perception on physical activity during hospitalization needs further attention.

### Factors influencing physical activity promotion

The identified factors influencing physical activity promotion by nurses, most importantly patient motivation and high workload causing priority shifts of tasks, are in line with previous studies addressing barriers in physical activity promotion during hospitalization (10, 13, 36, 37). The nurses in our study suggested to increase patient awareness on the importance of physical activity. Barriers for being physically active during the hospital admission from a patient perspective were previously addressed (10), but better understanding of what causes low patient motivation is important.

According to the nurses in our study, physical activity promotion tasks became less of a priority when nurses experienced a low staff ratio and a high workload. Staff ratio and workload are associated with nursing tasks being left undone which was found to be related to the nurses perception of quality of nursing care (38). In addition, an increase in nurses’ workload was found to affect patient outcomes (39). This implies that nurse staffing levels should be increased or tasks must shift towards other actors or targeted by eHealth interventions (40). However, intervention studies on physical activity promotion of hospitalized patients showed positive results on physical activity levels using preexisting staff ratios (41, 42).

In the current study, physical activity promotion by the physician and carers involvement were indicated as influencing factors. Physicians were expected to indicate the ability of patients to be physical active. Physicians orders regarding physical activity are found to influence patients’ decisions to perform physical activity during hospitalization (43) but are infrequently discussed (43) and often bed rest orders during hospital admission do not have valid and specified reasons (6). Awareness of the role of physicians in physical activity promotion might contribute to strengthen nurses’ physical activity promotion behavior and increase physical activity levels of older hospitalized patients. Furthermore, carers could play a more prominent role in physical activity promotion. In our study the role of carers in physical activity promotion during hospitalization was indicated as minor, but including carers in physical activity promotion of older patients was previously emphasized (13).

### Strengths and limitations

This study used a mixed-methods approach and included a large group of nurses to explore the tasks and responsibilities in physical activity promotion of older patients during hospitalization. The questionnaire and interview design were self-developed and not cross validated. The use of another validated instrument was not possible while there was none available for this specific interest. However, both questionnaire and interview design were based on literature and tested in advance on feasibility and interpretation of the questions.

## Conclusion

Nurses perceive to have various roles in physical activity promotion and feel responsible, but they were not satisfied with the level of physical activity of patients. Contributing factors were low patient motivation and priority shifts due to high workload. Hospital managers and health care professionals should be aware of the various roles of nurses in physical activity promotion. Emphasis should be on the multidisciplinary approach of physical activity promotion including physicians, patients, and carers.

## Acknowledgements

We thank the nurses for participating, Irene Jongerden for her help during interview design and Melina van Gunsteren for her advice during drafting the manuscript.

